# An anterior hypothalamic circuit gates stress vulnerability

**DOI:** 10.1101/2024.10.28.620614

**Authors:** Zachary T Pennington, Alexa R LaBanca, Shereen D Abdel-Raheim, Madeline E Bacon, Afra N Mahmoud, Patlapa Sompolpong, Austin M Baggetta, Yosif Zaki, BumJin Ko, Zhe Dong, Alexander CW Smith, Paul J Kenny, Denise J Cai

**Affiliations:** Nash Family Department of Neuroscience, Icahn School of Medicine at Mount Sinai; Department of Pharmacology, Icahn School of Medicine at Mount Sinai

## Abstract

Prior adversity increases susceptibility to subsequent stressful events, but the causal underlying changes in brain circuitry are poorly understood. We harnessed unbiased whole-brain activity mapping to identify circuits that are functionally remodeled by prior adversity. This revealed that the anterior hypothalamic nucleus (AHN) displays heightened stress reactivity in previously stressed mice. This was accompanied by increased functional connectivity between the AHN and a threat-related limbic network. Using *in vivo* Miniscope imaging, we found that neuronal activity in the AHN encodes stressor valence. Moreover, stimulating AHN neurons enhanced, and inhibiting their activity mitigated, reactivity to stressful events. Lastly, silencing amygdala inputs to the AHN abolished the ability of prior adversity to increase stress sensitivity. These findings define a key role of the AHN in gating stress vulnerability by scaling valence signals from the amygdala.

## Introduction

The brain’s response to stress is fundamentally protective, engaging physiological and behavioral adaptations that promote survival. However, stressful experiences can also precipitate maladaptive brain plasticity that increases susceptibility to conditions like post-traumatic stress disorder (PTSD), anxiety, depression, and substance use disorder (*1–5*). Notably, there is substantial variation in how individuals cope with stressful life events. Exemplifying this, only a small proportion of individuals who experience a traumatic event will at some point develop PTSD or other trauma-related illness (*3*). How variability in stress circuits in the brain confers susceptibility to trauma-related illnesses is poorly understood.

The prior experience of stress – whether in the form of early childhood adversity or adult psychological trauma – is one of the most reliable predictors of adverse reactions to subsequent stressful experiences (*1, 2, 6–8*). To date, research on stress vulnerability has predominantly focused on a small subset of brain regions, among them the amygdala, prefrontal cortex, nucleus accumbens, and hippocampus (*9–11*). These efforts have yielded important insights into the cellular, molecular, and circuit sequelae of stress (*11–14*). However, in order to discover how these canonical regions interact with broader brain circuits, and to identify novel targets for disease intervention, we must also pursue more exploratory approaches.

Here, we capitalized on an unbiased discovery-based approach to map brain-wide neuronal activity patterns in concert with a “two-hit” stress procedure that renders mice susceptible to stressful events. We identify the anterior hypothalamic nucleus (AHN) as a novel regulator of stress vulnerability and find that functional connectivity between the AHN and other stress-related brain regions is increased following stress. Further, we demonstrate that neurons within this under-studied region potently encode the valence of stressful events, and that their activity is able to bi-directionally regulate stress responses. Lastly, we demonstrate the AHN interacts with canonical stress circuits to scale the impact of stressful events. Accordingly, the AHN represents a compelling new target for understanding stress susceptibility.

## Results

### Prior stress enhances AHN response to a future stressor

To investigate how prior adversity influences brain-wide reactivity to subsequent stress, we leveraged our previous observation that mice subjected to a strong stressor show a long-lasting and experience-dependent sensitization of subsequent stress responses (Fig 1A) (*15*). Mice were subjected to a high-intensity stressor (Stressor 1), in which they received 10 footshocks (Stressed, S), or they were placed in the same environment but did not receive footshocks (Non-Stressed, NS). Then, ∼10 days later, both groups of mice were exposed to a loud auditory stressor while in their homecages (Stressor 2). Critically, animals that underwent Stressor 1 showed a heightened defensive freezing response to Stressor 2 (Fig 1B), resembling the sensitizing impact of prior adversity on stress responses in humans (*1, 2, 6–8*).

**Figure 1.**
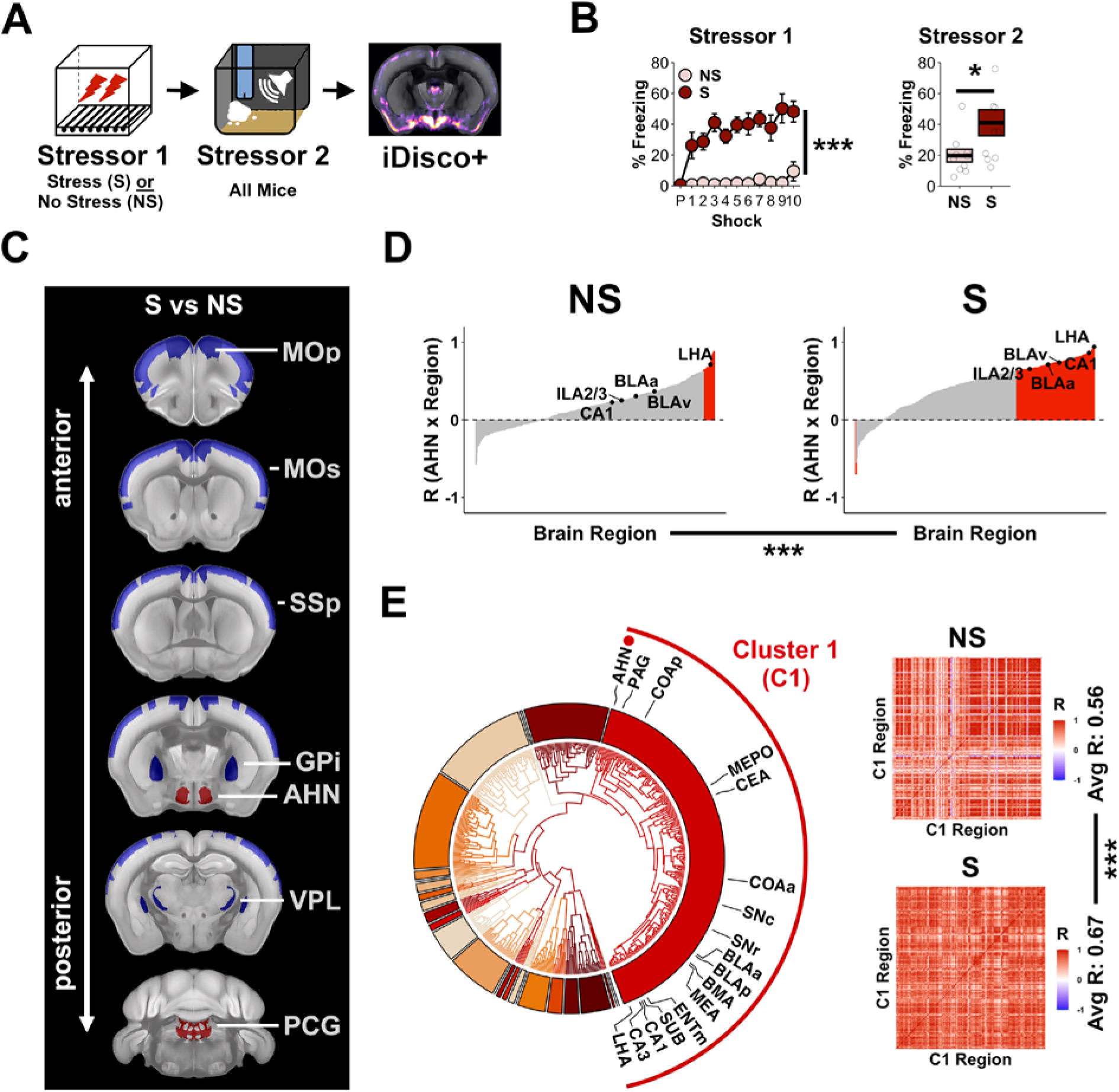
Prior stress increases activity and functional connectivity of the AHN. A) Mice underwent Stressor 1, consisting of 10 footshocks (S), or were placed in the same environment and received no stress (NS). 10 days later, all mice received Stressor 2, a loud auditory stimulus. Intact brains were then cleared and brain-wide cFos was quantified. N=10/group. B) Mice that received Stressor 1 (S) displayed increased freezing relative to mice that did not (NS), both during Stressor 1 (left), and during Stressor 2 (right). RM-ANOVA for Stressor 1 freezing – Group: F_1,18_=113.5, p<0.001. ANOVA for Stressor 2 freezing – Group: F_1,19_=4.6, p=0.046. C) Group differences in cFos aligned to the Allen Brain Atlas. Animals that received Stressor 1 (S) showed broad cortical hypoactivation coupled with subcortical hyperactivation of the AHN and PCG. Red = S>NS. Blue = S<NS. See Tables S1-S2 for statistics and abbreviations. D) Animals that received Stressor 1 (S) showed an increased number of correlations between the AHN and other brain regions, including several limbic brain regions. Chi-square contingency test of S vs NS: χ^2^=116.09, p<0.001. E) Left) Hierarchical clustering places the AHN in a cluster (Cluster 1) that includes many limbic brain regions. Right) Previously stressed animals (S) display higher intra-cluster correlational strength (right) (permutation test, p<0.001). p<.05 (*), p<0.01 (**), p<0.001 (***). Error bars reflect standard error of the mean.

To identify brain-wide activation patterns associated with this sensitized stress response, intact brains were stained for the activity-dependent immediate early gene cFos and cleared using the iDISCO+ method (*16*). Three-dimensional images were then acquired using light-sheet microscopy, processed, and aligned to the Allen Brain Atlas, permitting group differences in cFos activity across >450 regions to be assessed in an unbiased manner (Figs 1C, S1; Tables S1-S2).

Mice previously exposed to Stressor 1 showed broad cortical hypoactivation in response to Stressor 2 relative to mice not subjected to Stressor 1, particularly in superficial prefrontal and somatomotor cortical layers (Fig 1C; see Tables S1-2 for all statistics and abbreviations). This is consistent with previous reports showing hypoactivation of cortical regions in previously stressed mice (*17, 18*), and stress-induced alterations in cortical function (*19*). Additionally, previously stressed mice displayed hyperactivation of the pontine central gray (PCG), a region that has been shown to be critical for auditory startle responses (*20, 21*). They also displayed hyperactivation of the anterior hypothalamic nucleus (AHN), a brain region implicated in defensive behaviors but is relatively understudied (*22–25*). Critically, the AHN is well positioned to regulate stress susceptibility based on its dense connectivity with limbic and hypothalamic brain regions (*26*). Lastly, although several stress-associated regions did not show differential activation based upon stress history, we validated that these regions showed heightened cFos relative to control animals that did not receive Stressor 2 (Fig S2

### Prior stress remodels AHN functional connectivity

A major advantage of whole-brain cFos mapping is the ability to extend beyond group comparisons of discrete brain regions, to build models of functional connectivity across the brain. Given that the AHN is interconnected with established stress circuits (*26*), we investigated whether a prior stressful experience biases AHN connectivity with other brain regions in response to a subsequent stress. Examining the correlations between cFos expression in the AHN and every other mapped structure, previously stressed animals (S) demonstrated connectivity between the AHN and other stress-associated limbic regions, particularly the amygdala, hippocampus, and medial prefrontal cortex (Fig 1D). Much smaller numbers of correlations between the AHN and other brain regions were observed in the animals that did not experience Stressor 1 (NS, Fig 1D). This suggests that prior experience increases functional connectivity between the AHN and brain regions involved in processing stress.

To further explore the possibility that the AHN might be part of a brain network whose activity is modified by prior experience, we next performed hierarchical clustering of brain regions in the previously stressed mice. This approach allows for natural segregation of brain regions into clusters based upon their stress-evoked co-activity. Taking all region-region correlational pairs into account, we found that the AHN was situated in a large cluster of stress-related regions (Fig 1E, Cluster 1). Once again, this AHN cluster was composed of stress-related limbic regions including amygdala nuclei (COAp, CEA, BLAa, BLAp, BMA, MEA), hippocampal nuclei (CA1, CA3, SUB), the periaqueductal gray (PAG), the lateral hypothalamus (LH), and substantia nigra (SNr, SNc). Of relevance, several of these structures have been found to have monosynaptic connections with the AHN (*23, 24, 26*). Lastly, we assessed if prior stress might influence the coordinated activity of this cluster of AHN-related brain regions. We found that the average correlational strength of Cluster 1was substantially higher in previously stressed relative to unstressed animals (Fig 1E), and this change was not due to higher brain-wide correlational strength (Fig S3). These findings suggest that prior adversity increases the functional connectivity of the AHN with a network of threat-related brain regions.

### GABAergic AHN neuron activity reflects stressor valence

Immediate early gene imaging allowed us to identify the AHN as a putative regulator stress vulnerability, but lacks the temporal precision necessary to identify what features of stress, or of the stress response, precipitate changes in AHN activity. Moreover, few studies had recorded from AHN neurons *in vivo* to determine the features of stress encoded by these neurons. Therefore, we tracked the activity of individual AHN neurons, employing calcium imaging of freely behaving mice with open-source Miniscopes (Fig 2A-C) (*27*). GABAergic AHN neurons were recorded because they are the dominant neuronal population within the AHN (*23, 24*), and many GABAergic AHN neurons provide efferent synaptic input to downstream brain regions (*28*). We found that a large population (∼40%) of AHN neurons reliably respond to the onset of the footshock stress (Fig 2D). Moreover, the amplitude of AHN calcium transients was closely correlated with the intensity of the footshock (Fig 2D).

**Figure 2.**
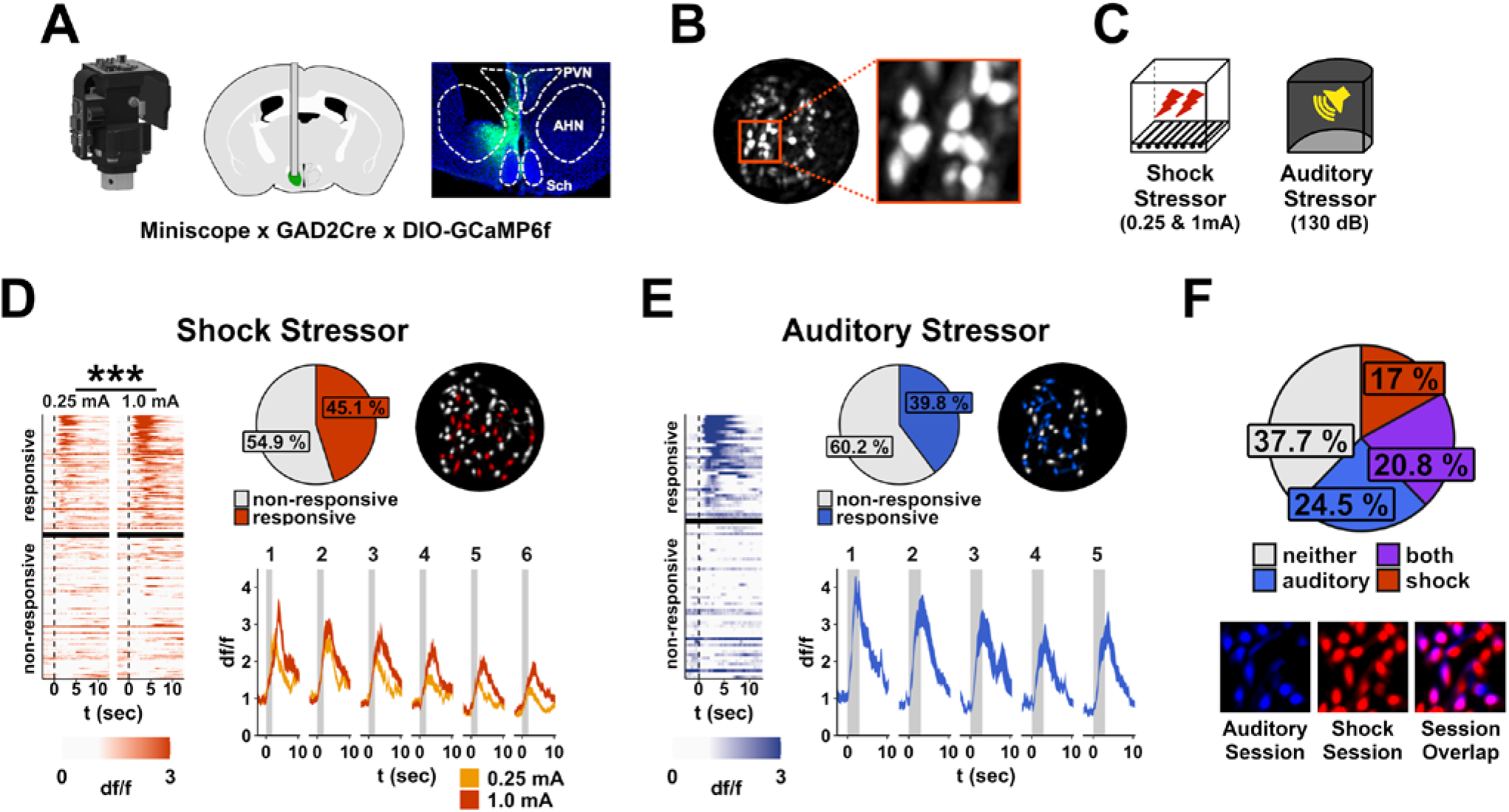
GABAergic AHN neurons reflect stressor valence. **A)** To track neuronal activity of the AHN in freely behaving mice, GCaMP6f was expressed in GABAergic AHN neurons and a GRIN lens was implanted overlying the AHN. Activity was recorded with a Miniscope. **B)** Example of a maximum projection across a Miniscope recording session showing the field of view. **C)** To examine AHN neuronal responses to stress, animals were exposed to low and high amplitude footshock, as well as to an auditory stressor. **D)** GABAergic neurons respond to footshock in a graded fashion. Left) Each row represents the average response of a neuron to low and high amplitude footshock. Responsive neurons show a stronger response to high (vs low) amplitude shock (RM-ANOVA for post-shock activity – Amplitude: F_1,73_=30.1, p<0.001) Top-middle) Proportion of neurons that reliably respond to footshock. Top-right) Example field of view, pseudo-colored to depict spatial location of shock-responsive cells in red. Bottom-right) Average activity of shock-responsive cells to low and high amplitude shocks across 6 trial pairs. **E)** GABAergic neurons respond to auditory stressor. Left) Each row represents the average response of a neuron to an auditory stressor. Top-middle) Proportion of neurons that reliably respond to auditory stressor. Top-right) Example field of view, pseudo-colored to depict spatial location of auditory-responsive cells in blue. Bottom-right) Average activity of responsive cells to auditory stressor across 5 trials. **F)** A proportion of GABAergic AHN neurons respond to both footshock and auditory stress. Top) Proportion of neurons that that respond to footshock stress, auditory stress, both stressors, or neither stressor. B) Example showing alignment of neurons across sessions. p<.05 (*), p<0.01 (**), p<0.001 (***). Error bars reflect standard error of the mean.

Then, to assess if AHN neurons respond specifically to footshock/somatosensory events, or if they might provide a broader signal about stressful events, we examined the response of AHN neurons to a stressful auditory stimulus. We found a similar proportion of neurons within the AHN reliably respond to this stimulus as to footshock (Fig 2E). In addition, to see if the same population of AHN neurons might encode different stressor modalities, we cross-registered cells across footshock and auditory stressor sessions (Fig 2F). Although a fraction of neurons responded uniquely to either footshock (∼17%) or auditory stress (∼24.5%), a nearly equivalent proportion of neurons responded to both stimuli (20.8%). These findings indicate that at least a subpopulation of GABAergic AHN neurons respond to multiple aversive stimuli and dynamically track the intensity of the stressor (i.e., stressor valence).

### Augmenting AHN activity amplifies stressor valence

The preceding results suggest that the AHN encodes stressor valence and that prior stress might alter this encoding, sensitizing the behavioral response to stressful stimuli. To directly test the possibility that facilitation of AHN activity by prior stress augments stressor valence, we expressed the excitatory chemogenetic receptor HM3Dq in GABAergic AHN neurons, or a control virus expressing mCherry (Figs 3A, S4). Applying the agonist clozapine-*N*-oxide (CNO) to excite these neurons (Fig 3B), we hypothesized that this manipulation would promote defensive responding to an auditory stressor. As anticipated, activation of AHN neurons increased post-stress freezing (Fig 3C).

**Figure 3.**
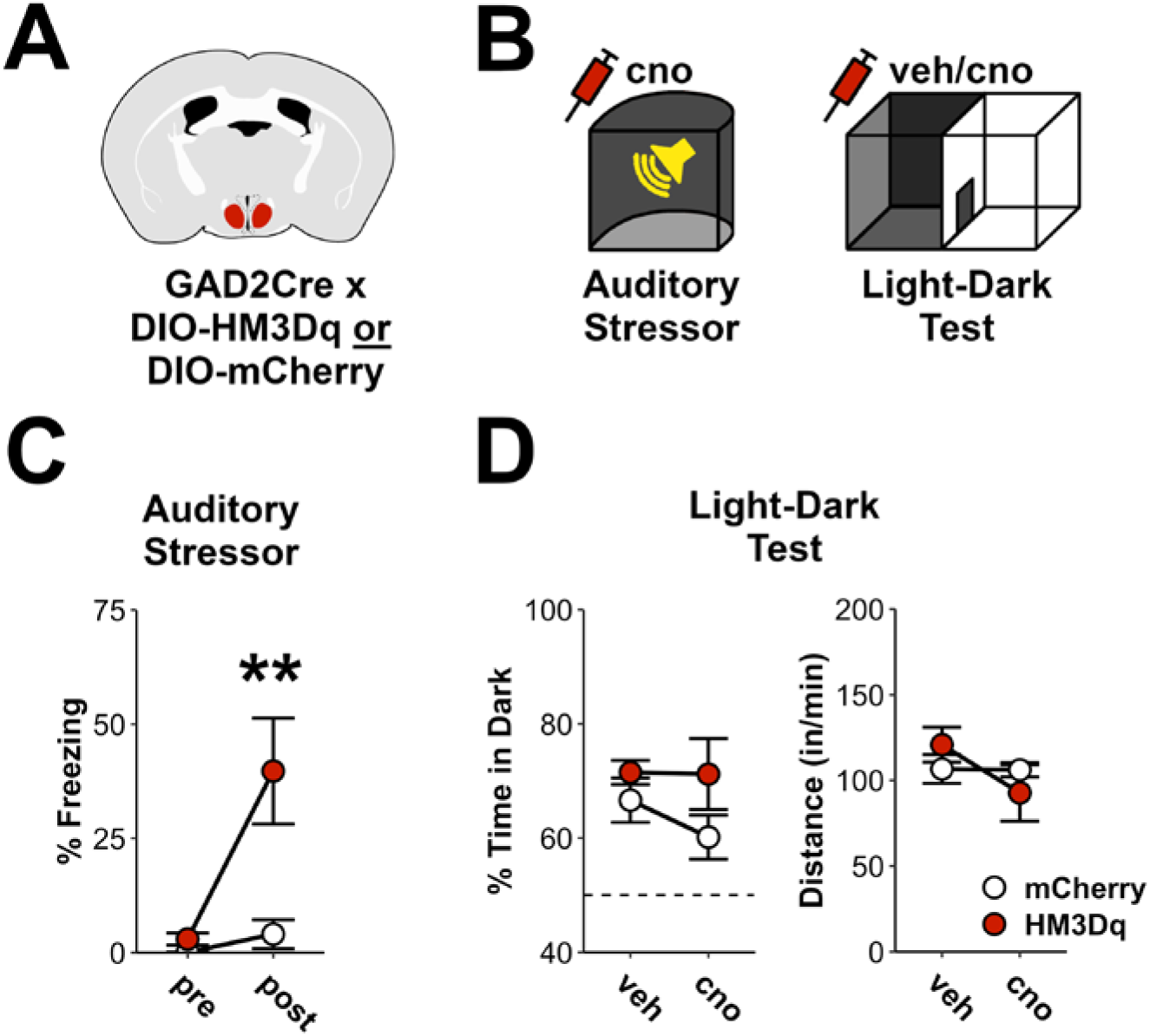
Augmenting AHN activity amplifies stressor valence. **A)** To excite GABAergic AHN neurons, a cre-dependent virus expressing HM3Dq, or a control virus expressing mCherry, was infused into the AHN of GAD2Cre mice. N = 9 HM3Dq (4 female), 12 mCherry (6 female). **B)** The agonist cno was administered before delivery of an auditory stressor. Additionally, mice were tested in the light-dark test twice, on and off cno (order counter-balanced). **C)** Activation of GABAergic AHN neurons had no effect on baseline freezing, but increased freezing after an auditory stressor. RM-ANOVA for freezing – Virus x Time: F_1,19_=10.9, p<0.01. Pre – Group: F_1,19_=4.2, p=0.053. Post – Group: F_1,19_=7.8, p=0.01. **D)** Activation of GABAergic AHN neurons had no effect on the proportion of time animals spent in the dark, nor on locomotor activity. RM-ANOVA for time in dark – Virus: F_1,19_=2.7, p=0.12; Virus x drug: F_1,19_=0.8, p=0.39. RM-ANOVA for distance travelled – Virus: F_1,19_=0, p=0.96; Virus x drug: F_1,19_=1.4, p=0.25. p<.05 (*), p<0.01 (**), p<0.001 (***). Error bars reflect standard error of the mean.

The facilitation of stress-elicited freezing by AHN stimulation could reflect an amplification of the auditory stressor’s valence. Alternatively, it may be that activation of the AHN is itself stressful or promotes a general anxiety-like state. First, pre-stressor freezing was not altered by AHN activation (Fig 3C), suggesting this manipulation is not aversive. Second, testing the same animals in the light-dark test to assess anxiety-related behavior (*29*), we found that activating AHN neurons did not influence time spent in the dark, the primary metric of anxiety-related behavior, nor did it alter locomotion (Fig 3D). Thus, enhanced neural activity in the AHN appears to amplify the valence of stressful stimuli.

### AHN activity is necessary for the induction and expression of stress-induced defensive behavioral changes

Having found that AHN activity is sufficient to increase stress reactivity, we next asked if AHN neuronal activity is necessary for stress-induced changes in defensive behavior (i.e., threat-elicited behavior) (Fig 4). We first tested the effect of silencing AHN neurons during a strong footshock stressor (Stressor 1), utilizing the inhibitory opsin stGtACR1 (GtACR, Figs 4A-F, S5) (*30*). Acute inhibition of AHN neurons drastically reduced freezing during Stressor 1 (Fig 4C), consistent with a reduction in stressor valence. Moreover, testing these animals days later when the AHN was no longer inhibited, we found that prior AHN inhibition during Stressor 1 reduced subsequent freezing in the shock-associated context (Stressor 1 Recall, Fig 4D). Additionally, AHN inhibition during Stressor 1 reduced the response to a subsequent auditory stressor (Stressor 2, Fig 4E,F). This shows that AHN activity is necessary not only for the immediate defensive freezing response to footshock, but the induction of persistent defensive behavioral changes afterward, such as recall of the stress memory (Stressor 1) and sensitization to subsequent stress (Stressor 2). Importantly, we have previously found that reducing stressor valence through a reduction in shock amplitude reduces these same behaviors (*15*).

**Figure 4.**
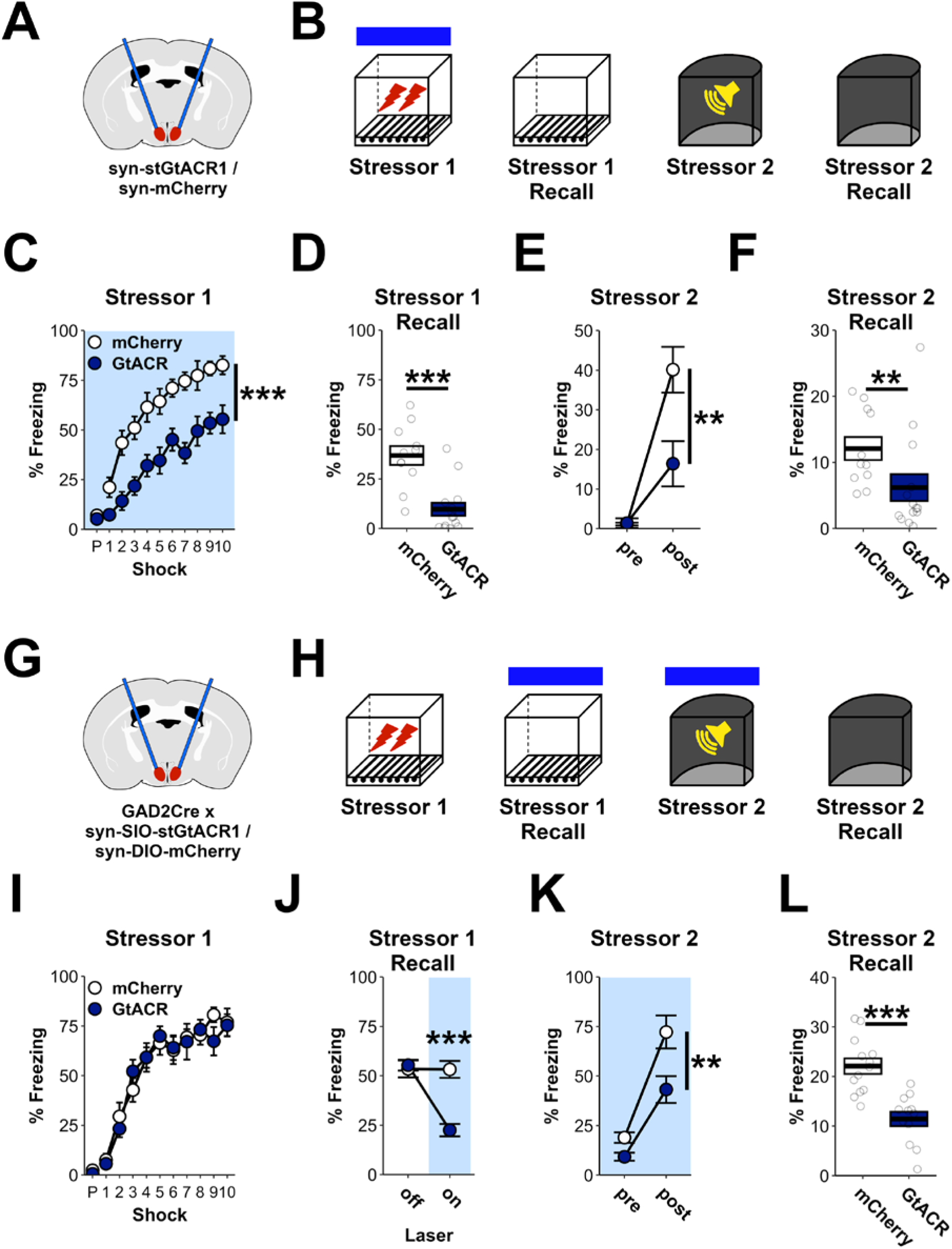
AHN activity is necessary for the induction and expression of stress-induced defensive behavioral changes. **A)** To pan-neuronally inhibit AHN neurons, a virus expressing the inhibitory opsin stGtACR1 (GtACR) or a control virus expressing mCherry was infused into the AHN. Optic fibers were implanted just overlying the AHN. N = 11 mCherry, 14 GtACR. **B)** Animals underwent a strong footshock stressor (Stressor 1) with light inhibiting AHN neurons throughout the session. With the AHN no longer inhibited, animals were then tested for their memory of the footshock context (Stressor 1 Recall), their response to an auditory stressor in a new context (Stressor 2), and their memory for that context (Stressor 2 Recall). **C)** Inhibition of AHN neurons potently reduced post-shock defensive freezing during Stressor 1. RM-ANOVA for freezing – Virus: F_1,25_=37.6, p<0.001, Virus x Shock: F_9,225_=0.8, p=0.51. **D)** Prior inhibition of AHN neurons during Stressor 1 reduced freezing during Stressor 1 Recall. ANOVA for freezing – Virus: F_1,25_=24, p<0.001. **E)** Prior inhibition of AHN neurons during Stressor 1 reduced freezing after Stressor 2. RM-ANOVA for freezing – Virus x time: F_1,25_=10.8, p<0.01. ANOVA for post-stressor freezing – Virus: F_1,25_=10.7, p<0.01. ANOVA for pre-stressor freezing – Virus: F_1,25_=0, p=0.96. **F)** Prior inhibition of AHN neurons during Stressor 1 reduced freezing during Stressor 2 Recall. ANOVA for freezing – Virus: F_1,25_=6.5, p=0.017. **G)** To inhibit GABAergic AHN neurons, a cre-dependent virus expressing the inhibitory opsin stGtACR (GtACR) or a control virus expressing mCherry was infused into the AHN of GAD2Cre mice. Optic fibers were implanted just overlying the AHN. N = 13 mCherry (6F), 12 GtACR (7F). **H)** AHN neurons were inhibited during Stressor 1 Recall to assess their contribution to fear memory recall, and Stressor 2, to assess their contribution to sensitized stress responses. **I)** No difference was observed between groups during Stressor 1, when the AHN was not inhibited. RM-ANOVA for post-shock freezing – Virus: F_1,23_=0, p=0.85; Virus x Shock: F_9,207_=0.7, p=0.73. **J)** Inhibiting GABAergic AHN neurons reduced freezing during Stressor 1 Recall. RM-ANOVA for freezing – Virus x Light: F_1,23_=54, p<0.001. **K)** Inhibiting GABAergic AHN neurons reduced freezing during Stressor 2. RM-ANOVA for freezing – Virus: F_1,23_=12.4, p<0.01; Virus x Time: F_1,23_=2.7, p=0.11. **L)** Prior inhibition of GABAergic AHN neurons during Stressor 2 reduced subsequent freezing during Stressor 2 Recall. ANOVA for freezing – Virus: F_1,23_=23.2, p<0.001 p<.05 (*), p<0.01 (**), p<0.001 (***). Error bars reflect standard error of the mean.

Next, we tested the necessity of GABAergic AHN neurons for the expression of stress-induced changes in defensive behavior (Fig 4G-L). After experiencing an initial footshock stressor (Stressor 1, Fig 4I), silencing GABAergic AHN neurons reduced freezing when animals were returned to the stressor context (Stressor 1 Recall, Fig 4J). This indicates that these neurons are necessary for responding to stress-associated cues, in addition to responding to stressors themselves. Moreover, similar to the effect of silencing AHN neurons during Stressor 1, silencing these neurons during a subsequent auditory stressor (Stressor 2) blunted freezing across the session (Fig 4K), as well as when animals were placed back into this context when these neurons were no longer inhibited (Stressor 2 Recall, Fig 4L). Taken together, these results demonstrate that the AHN is necessary for both the induction and expression of stress-induced defensive behavioral changes.

### An amygdala-hypothalamic circuit gates stress vulnerability

Finally, we investigated how AHN neurons respond to inputs from upstream brain regions to regulate behavioral responding to stressful events (Fig 5). The AHN is known to receive input from stress-associated regions, among them the basolateral amygdala (BLA) and the ventral hippocampus (vHC) (Fig S6). Of interest, we have previously shown that stress-induced protein synthesis and subsequent neuronal activity in the BLA, but not the vHC, is necessary for stress-induced enhancements in negative valence (*15*). Thus, we speculated that neuronal projections from the amygdala to the AHN, but not the vHC, support heightened representations of stressor valence in animals that experienced prior adversity. To test this hypothesis, a retrograde virus expressing cre-recombinase was infused into the AHN and a cre-dependent virus expressing the inhibitory chemogenetic receptor HM4D was infused into either the BLA or vHC. Alternatively, a control virus was infused into these structures (Fig 5A). This allowed us to selectively silence BLA or vHC cells that directly project to the AHN (Fig 5B). After receiving an initial stressor (Stressor 1), CNO or vehicle was administered prior to recall of the initial stressor context (Stressor 1 Recall). Here, neither inhibition of BLA-AHN nor vHC-AHN projecting neurons was able to reduce freezing (Fig 5D). However, when CNO was administered prior to the second auditory stressor (Stressor 2), inhibition of BLA-AHN projecting neurons inhibited freezing, whereas inhibiting vHC-AHN projections was without effect (Fig 5E-F). These results are consistent with the established role of the amygdala in processing the valence of threatening stimuli (*31–33*), and suggest that the AHN gates valence signals from the BLA to modulate behavioral responses to stressful events.

**Figure 5.**
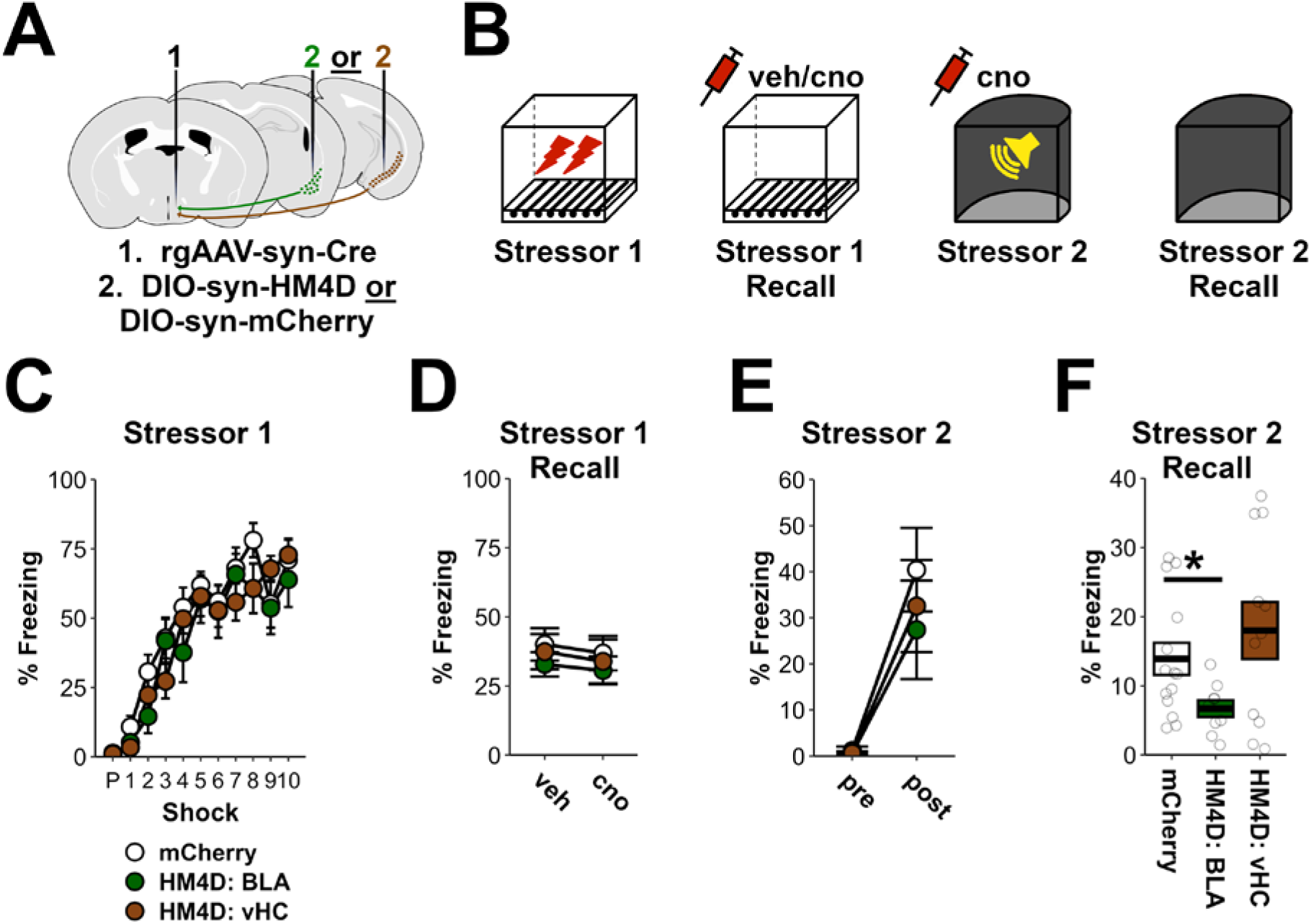
An amygdala-hypothalamic circuit gates stress vulnerability. **A)** To silence AHN inputs, a retrograde virus expressing cre-recombinase was infused into the AHN in combination with a cre-dependent HM4D-expressing virus, or a mCherry-expressing virus, in the BLA or vHC. N = 14 mCherry, 9 HM4D: BLA, 11 HM4D: vHC. **B)** AHN inputs were inhibited during Stressor 1 Recall to assess their contribution to fear memory recall, and during Stressor 2, to assess their contribution to sensitized stress responses. **C)** No difference was observed between groups during Stressor 1, when AHN inputs were not inhibited. RM-ANOVA for post-shock freezing – Group: F_2,31_=0.9, p=0.44; Group x Shock: F_18,279_=1, p=0.42. **D)** Inhibition of BLA and vHC inputs to the AHN had no impact on freezing during Stressor 1 Recall. RM-ANOVA for freezing – Group x Drug: F_2,31_=0, p=0.99. **E)** Inhibition of BLA and vHC inputs to the AHN had no impact on freezing during Stressor 2. RM-ANOVA for freezing – Group: F_2,31_=0.4, p=0.63; Group x Time: F_2,31_=0.4, p=0.66. **F)** Inhibition of AHN inputs from the BLA, but not the vHC, during Stressor 2, reduced freezing during Stressor 2 Recall. ANOVA for freezing – Group: F_2,31_=5.8, p<0.01; mCherry vs BLA: F_1,21_=6.9, p=0.02; mCherry vs vHC: F_1,23_=0.7, p=0.42. p<.05 (*), p<0.01 (**), p<0.001 (***). Error bars reflect standard error of the mean.

Finally, although inhibiting vHC inputs to the AHN had no effect on freezing during Stressor 1 or Stressor 2 recall, we found that silencing these neurons reduced stress-induced changes in anxiety-related behavior (Fig S7), in line with prior work on the vHC in anxiety-related behavior (*15, 34*). Additionally, we found that inputs from the lateral septum (LS) to the AHN regulate freezing during Stressor 1 Recall (Fig S7). Accordingly, the AHN appears to integrate inputs from diverse sources to regulate a range of defensive behaviors.

## Discussion

Above, we have identified the AHN as a putative regulator of stress vulnerability and its hub-like influence over stress-associated behavior. Stress-induced AHN activity was potentiated by the prior experience of stress, as was its connection with a threat-associated brain network. Additionally, recording from GABAergic AHN neurons *in vivo*, we found their activity reflected stress severity, suggesting they encode stressor valence. Providing causal support for this notion, manipulations of GABAergic AHN neurons were able to modulate behavioral responding to stressful stimuli in manner consistent with altering stressor valence. Lastly, AHN inputs from the amygdala – a region known for encoding valence – were found to similarly influence behavioral responding to aversive stimuli. These findings lead to the hypothesis that the AHN gates valence signals, and suggest that prior adversity may amplify AHN encoding of negative valence, resulting in heightened stress vulnerability.

This work elevates the need to complement investigations into known regions of interest with exploratory approaches in order to identify novel circuit interactions relevant to mental health. Prior work on the hypothalamus’ role in stress has most heavily focused on the paraventricular nucleus (PVN), a nucleus known to control release of the stress hormone cortisol via the hypothalamic-pituitary-adrenal axis (*35*). Notably, although the PVN receives diverse inputs (*36*), the AHN projects directly to the PVN, and in this way can directly modulate stress hormone release (*22*). That said, beyond its neuroendocrine actions, the AHN is likely to regulate stress-evoked behavior through complex extrahypothalamic interactions. These include monosynaptic projections to the LS, the PAG, and the amygdala (*37*), regions known to regulate behavioral responses to stress (*22, 38*). Indeed, a recent report found that AHN projections to the PAG are able to control attack behavior in response to painful stimuli (*39*). Combined with our observation that different inputs to the AHN are able to regulate distinct components of defensive behavior, the AHN appears to be a central hub regulating multiple defensive behaviors. While we have identified the critical role of amygdala inputs to the AHN in responding to aversive stimuli, future work is needed to more fully disentangle the complex input/output functions of the AHN, as well as how these interactions are modified by prior adversity. By understanding these relationships, novel targets for disease intervention might be discovered.

## ACKNOWLEDGEMENTS

This work was supported by NIMH DP2 MH122399, NIMH R01 MH120162, NIMH R56MH132959, Brain Research Foundation Award, Klingenstein-Simons Fellowship, NARSAD Young Investigator Award, McKnight Memory and Cognitive Disorder Award, One Mind-Otsuka Rising Star Research Award, Hirschl/Weill-Caulier Award, McKnight Brain Research Foundation & American Foundation for Aging Research Innovator Awards in Cognitive Aging and Memory Loss, and Chan Zuckerberg Initiative to DJC; NIMH K99 MH131792, BBRF Young Investigator Award, and Mount Sinai Friedman Brain Institute Postdoc Innovator Award to ZTP; The authors would like to thank Dr. Scott Russo, Dr. Roger Clem, and the members of the Cai and Shuman labs for their helpful comments on this work.

## AUTHOR CONTRIBUTIONS

ZTP and DJC conceived of the overarching research goals, designed the experiments, and oversaw the experiments. ZTP analyzed the experimental data and prepared the initial manuscript. ZTP, ARL, SDA, MEB, ANM, PS, AMB, YZ, BK, ZK, ACWS, PJK, and DJC contributed to interpretation of the results and edited the manuscript. ZTP, ARL, SDA, MEB, ANM, PS, AMB, YZ, BK, ZK, and ACWS performed experiments. ZTP and ZD designed software for analysis of behavioral data. DJC and ZTP secured funding.

## LIST OF SUPPLEMENTARY MATERIALS

Figures S1 to S4

Tables S1 and S2

Materials and Methods

## SUPPLEMENTARY MATERIALS FOR

**Figure S1.**
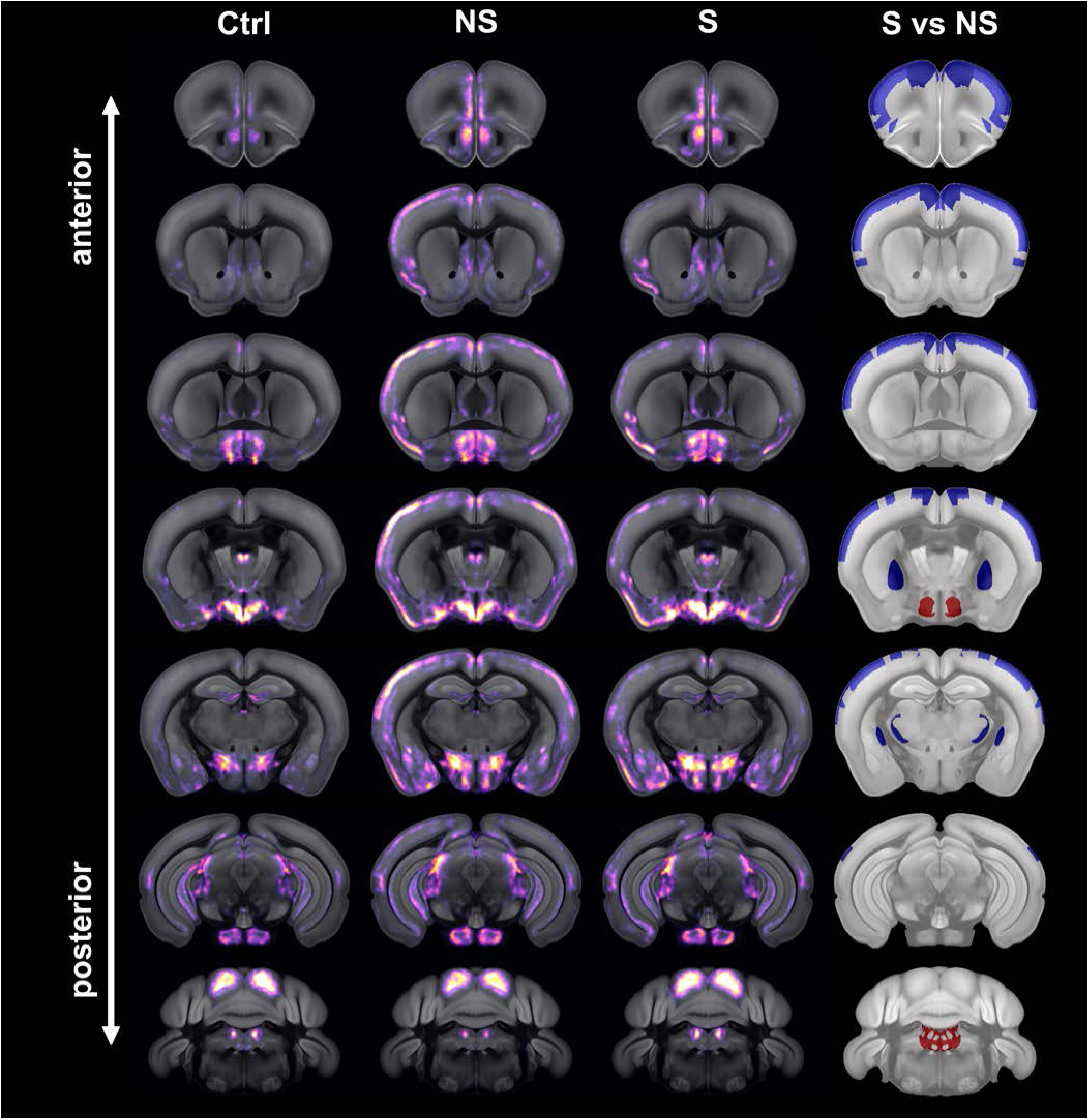
Distribution of whole-brain cFos activation across groups (Main Figure 1) Depiction of average cFos activation in control animals that did not receive Stressor 2 (Ctrl, 1^st^ column), animals that received Stressor 2 but not Stressor 1 (NS, 2^nd^ column), and animals that received both Stressor 1 and Stressor 2 (S, 3^rd^ column). Additionally, the contrast between groups S and NS is shown. Red = S>NS. Blue = S<NS. See Table S1 for statistics.

**Figure S2.**
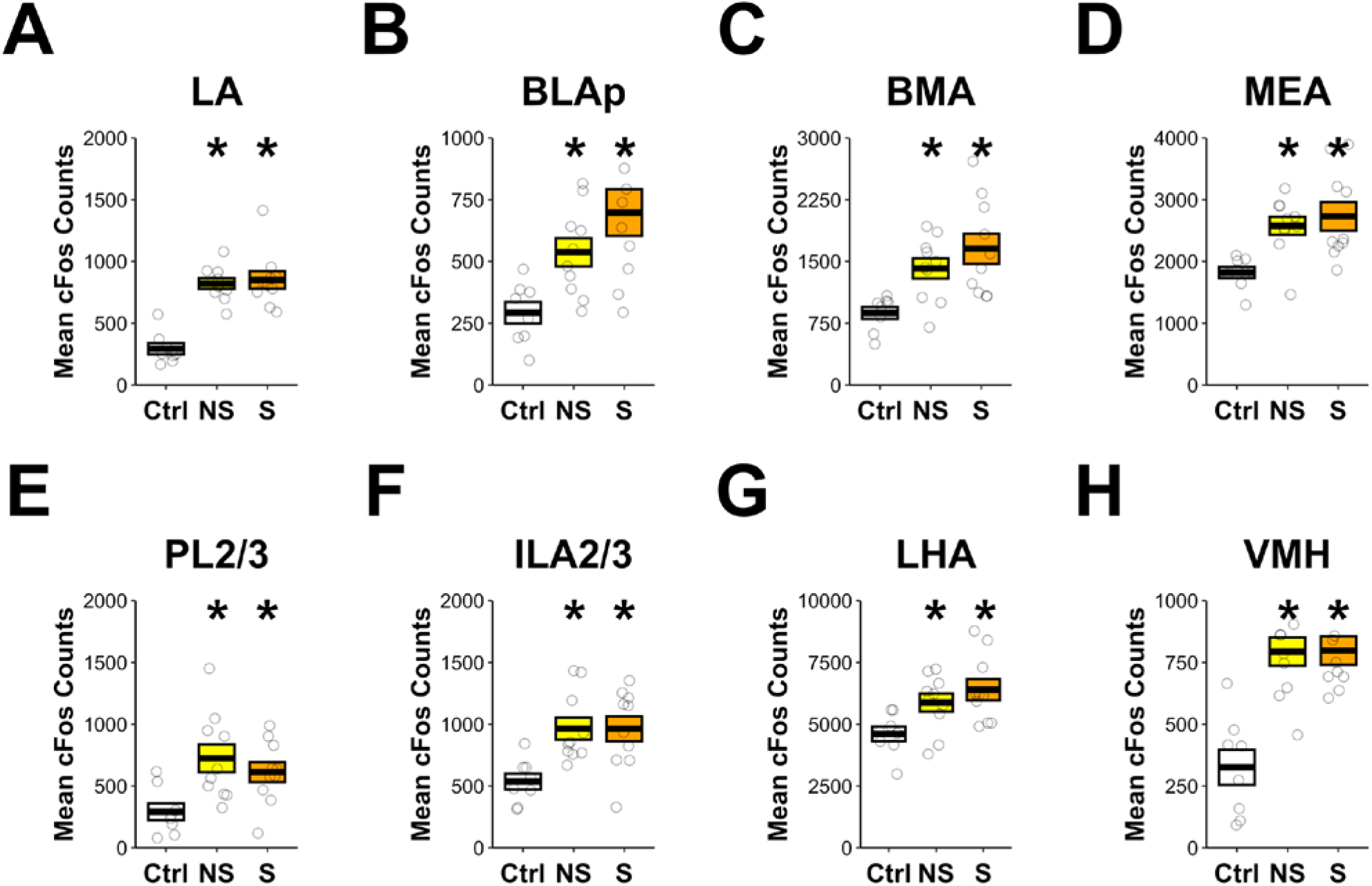
Stressor 2 activates a numerous brain regions associated with stress (Main Figure 1). Relative to animals that did not receive Stressor 2, animals that previously experienced Stressor 1 (S), as well as animals that did not (NS), display increased cFos in threat-related regions. This includes several amygdala subregions (LA, BLAp, BMA, MEA), prefrontal regions (PL2/3, IL2/3), and hypothalamic regions (LHA, VMH). * reflects difference from Ctrl. Boxplot represents mean and standard error. N = 8 Ctrl, 10 NS, 10 S. See Table S1 for statistics. LA = lateral amygdala; BLAp = posterior basolateral amygdala; BMA = basomedial amygdala; MEA = medial amygdala; PL2/3 = prelimbic, layers 2/3; ILA2/3 = infralimbic, layers 2/3; LHA = lateral hypothalamic area; VMH = ventromedial hypothalamus.

**Figure S3.**
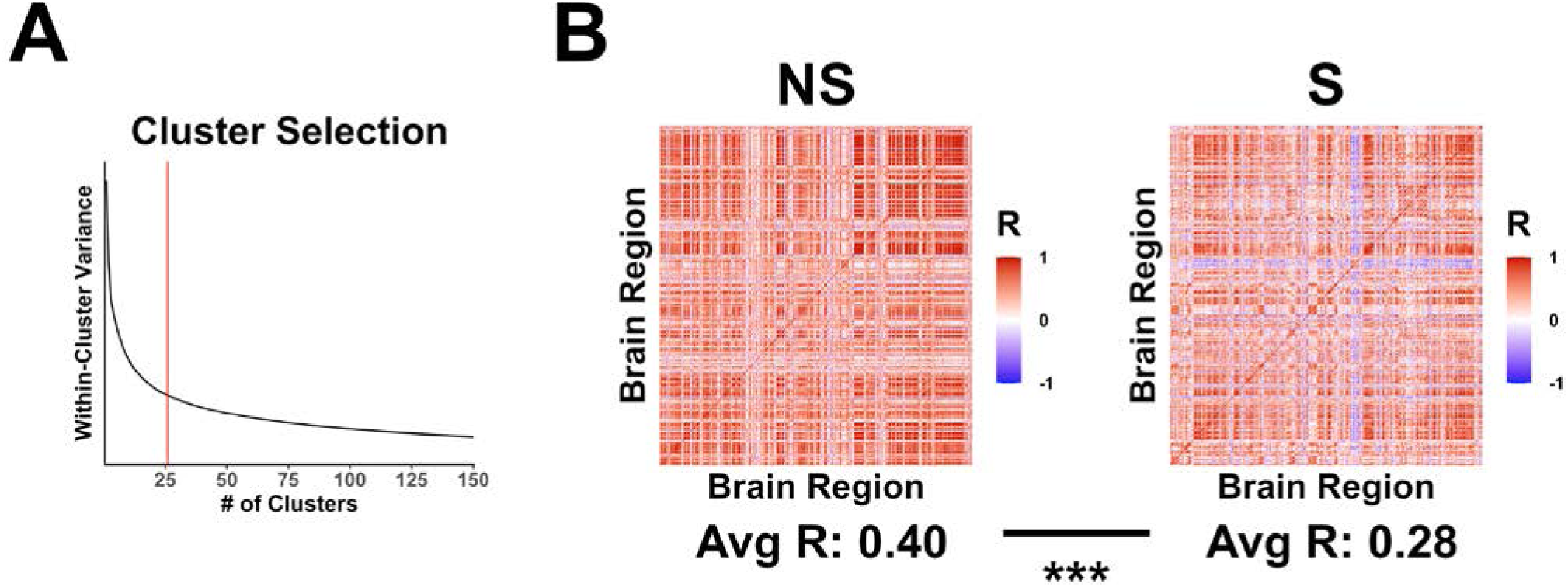
Selection of cluster cutoff and brain-wide correlation differences between groups (Main Figure 1). A) To select number of clusters for hierarchical clustering, we utilized the ‘elbow method’, in which the relationship between within-cluster variance and the number of clusters is plotted. The point at which within-cluster variance begins to stabilize is selected. Additionally, we attempted to keep average within-cluster correlations high (average R>0.5). B) Increased correlational strength within Cluster 1 for animals that previously received Stressor 1 (S) relative to those that did not (NS) could reflect a broad and non-specific increase in region-region correlational strength across the brain. Negating this possibility, it was actually found that when every brain region is examined, NS animals actually display higher region-region correlations on average. (permutation test, p<0.001). p<.05 (*), p<0.01 (**), p<0.001 (***).

**Figure S4.**
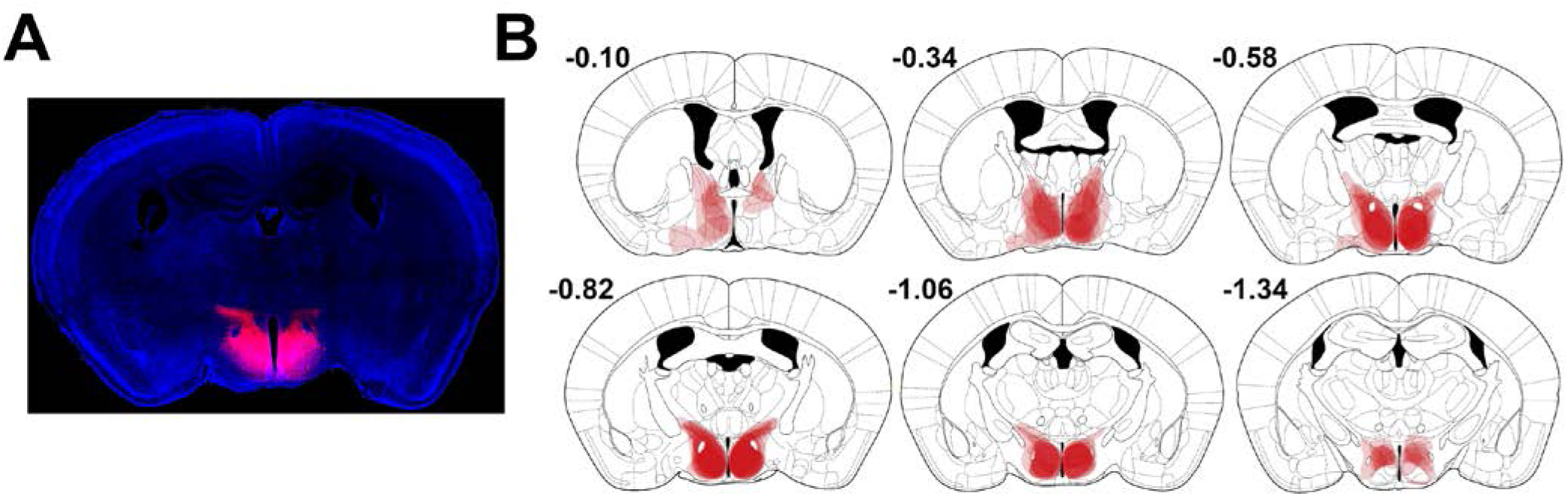
Distribution of HM3Dq in AHN (Main Figure 3). Viral placement of HM3Dq in AHN. A) Example of expression at center of injection site. B) Anterior-posterior distribution of expression. Expression was centered in the AHN, covering anterior, central and posterior components. There was occasional spread anterior into the medial preoptic nucleus, posteriorly into the dorsomedial hypothalamus, and dorsal into thalamus. Numbers adjacent to each coronal section correspond to anterior-posterior distance from bregma, in mm, according to the atlas of Franklin and Paxinos (*40*).

**Figure S5.**
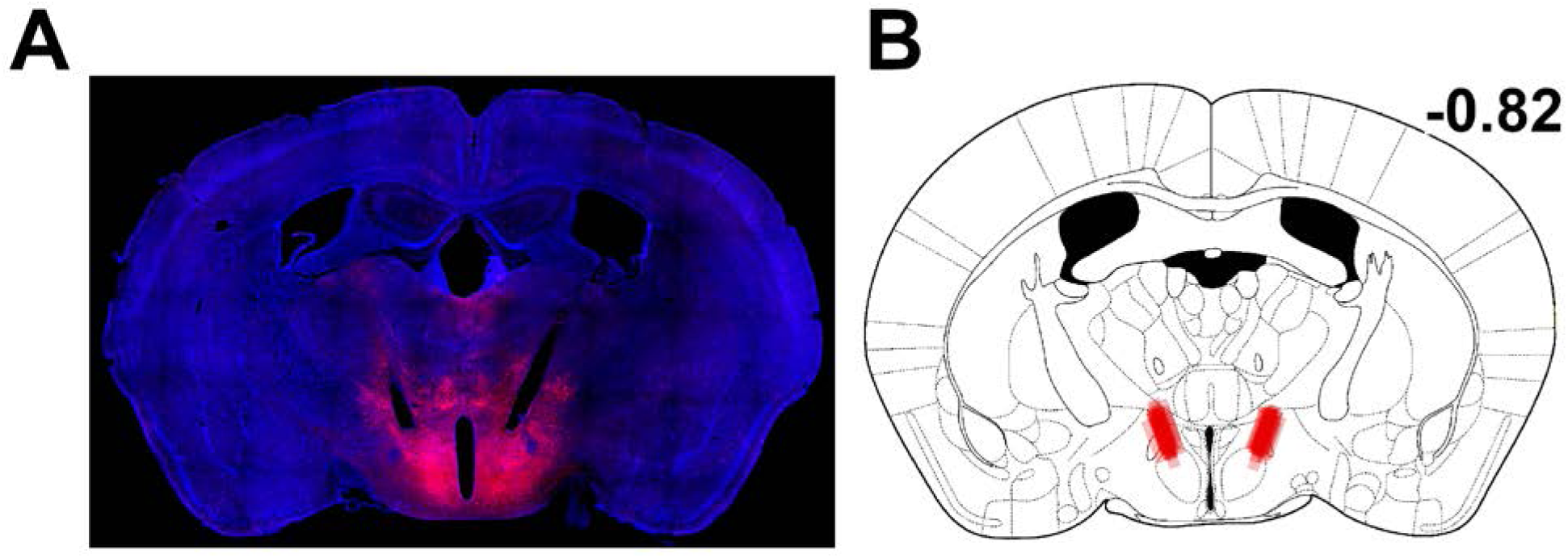
Distribution of optic fiber placement in AHN (Main Figure 4). A) Example of stGtACR expression at center of injection site. B) Distribution of fiber tip placement for all animals from the experiment presented in Fig 4A-F, representative of all optogenetic experiments. Fiber tips were located immediately above the central compartment of the AHN. Number adjacent to coronal section correspond to anterior-posterior distance from bregma, in mm, according to the atlas of Franklin and Paxinos (*40*).

**Figure S6.**
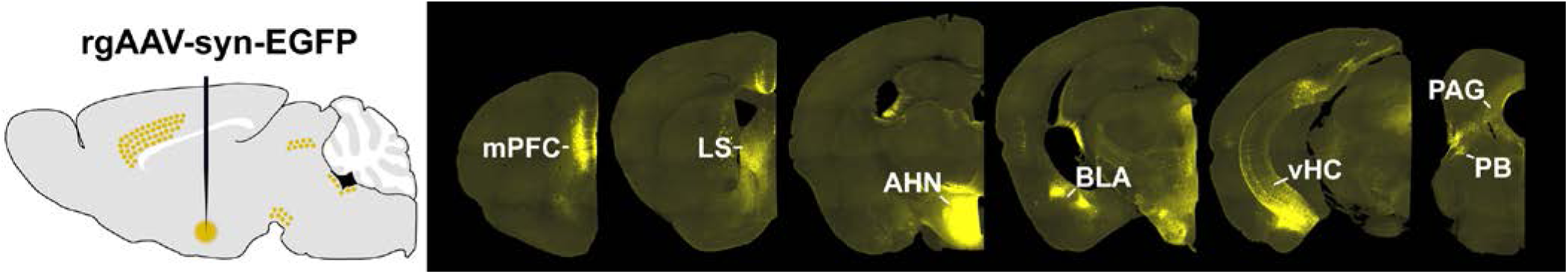
Retrograde tracing of inputs to AHN (Main Figure 5) Previous reports indicate that the AHN is innervated by several stress-related regions (*23, 24, 26*). Validating these findings, we infused a retrograde virus that expresses EGFP into the AHN (left). Consistent with prior reports, we found robust labeling of cells in the medial prefrontal cortex (mPFC), lateral septum (LS), basolateral amygdala (BLA), ventral hippocampus (vHC), periaqueductal gray (PAG), and parabrachial nuclei (PB).

**Figure S7.**
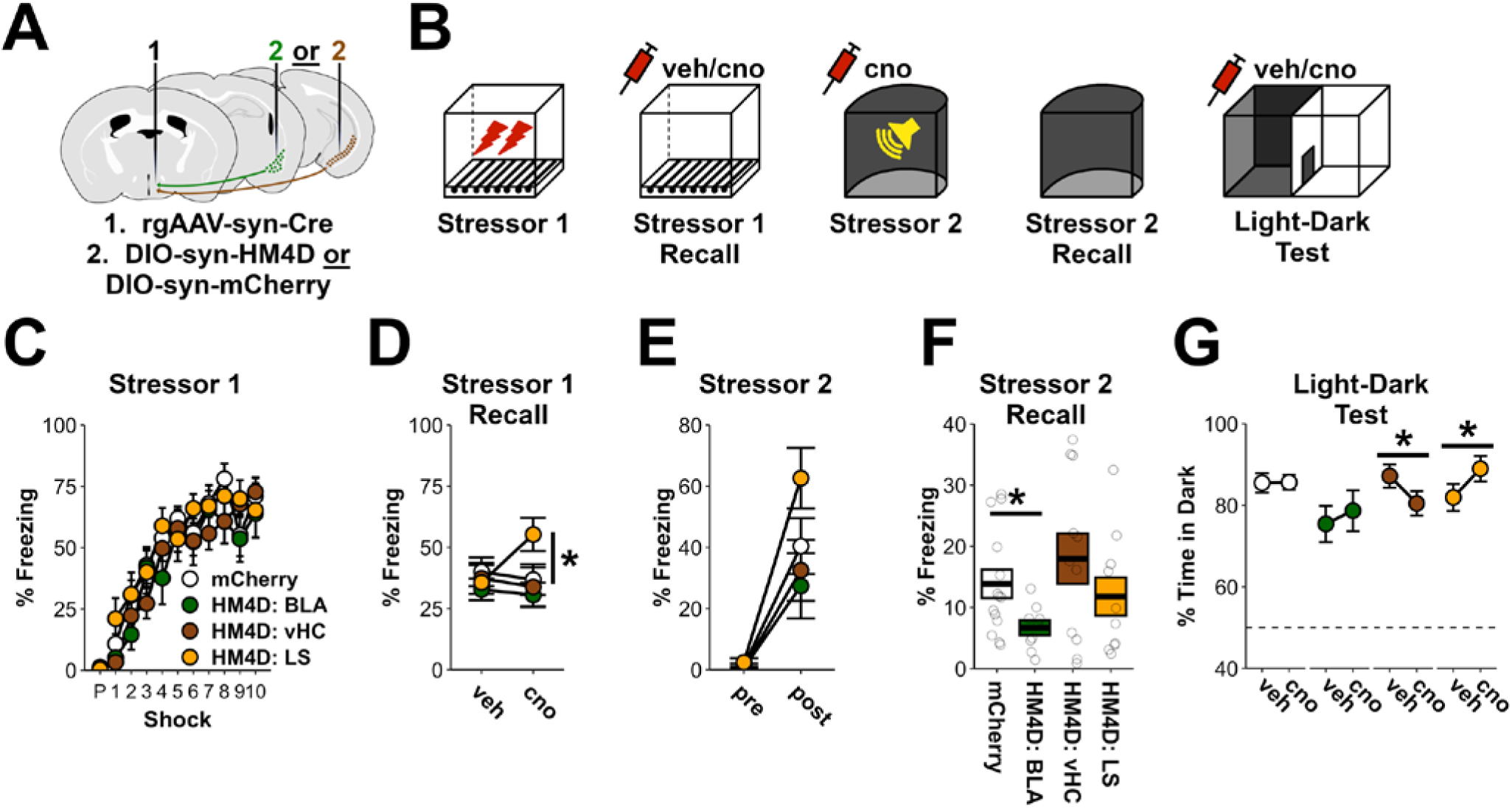
AHN regulates defensive behavior in an input-specific manner. **A)** To silence AHN inputs, a retrograde virus expressing cre-recombinase was infused into the AHN in combination with a cre-dependent HM4D-expressing virus, or a mCherry-expressing virus, in the BLA, vHC, or lateral septum (LS). Note that this is the same experiment from Main Figure 5 with data for LS group included, as well as a final set of light-dark tests. N = 14 mCherry, 9 HM4D: BLA, 11 HM4D: vHC, 10 HM4D: LS. **B)** AHN inputs were inhibited during Stressor 1 Recall to assess their contribution to fear memory recall; during Stressor 2, to assess their contribution to sensitized stress responses; and during the light-dark test, to assess their contribution to anxiety-related behavior. **C)** No difference was observed between groups during Stressor 1, when AHN inputs were not inhibited. RM-ANOVA for post-shock freezing – Group: F_3,40_=0.8, p=0.52; Group x Shock: F_27,360_=1, p=0.43. **D)** Inhibition of BLA and vHC inputs to the AHN had no impact on freezing during Stressor 1 Recall. However, inhibition of inputs from the LS to the AHN increased freezing. Given the predominantly inhibitory influence of LS outputs (*22*), this is consistent with a disinhibitory effect on the AHN. RM-ANOVA for freezing – Group x Drug: F_3,40_=2.34, p=0.09. Post-hoc test LS – Drug: F_1,9_=9.7, p=0.01. **E)** Inhibition of BLA and vHC inputs to the AHN had no impact on freezing during Stressor 2. RM-ANOVA for freezing – Group: F_3,40_=2.4, p=0.08; Group x Time: F_3,40_=2, p=0.13. **F)** Inhibition of AHN inputs from the BLA, but not the vHC, during Stressor 2, reduced freezing during Stressor 2 Recall. ANOVA for freezing – Group: F_3,40_=4.1, p=0.01; mCherry vs BLA: F_1,21_=6.9, p=0.02; mCherry vs vHC: F_1,23_=0.7, p=0.42; mCherry vs LS: F_1,22_=0.3, p=0.62. **G)** Inhibition of inputs to the AHN influenced anxiety-related behavior. Whereas inhibiting vHC inputs reduced time spent in the dark, consistent with prior reports inhibiting vHC (*15, 41*), inhibiting LS inputs increased time spent in the dark. ANOVA for time in dark – Group x Drug: F_3,40_=3.9, p=0.02. Post-hoc test of effects of drug on time in dark – BLA: F_1,8_=0.9, p=0.38; vHC: F_1,10_=5.6, p=0.04, LS: F_1,9_=7.7, p=0.02; mCherry: F_1,13_=2.2, p=0.96. p<.05 (*), p<0.01 (**), p<0.001 (***). Error bars reflect standard error of the mean.

## MATERIALS AND METHODS

### ANIMALS

All mice were bred on a C57BL/6J genetic background and were approximately 2-6 months old at the start of testing. Male C57BL/6J were obtained from Jackson Laboratories for whole-brain immediate early gene imaging as well as pan-neuronal optogenetic silencing (Jackson Laboratories, #000664). Studies with homozygous GAD2-IRES-Cre mice (Jackson Laboratories, #028867) utilized an equal mixture of female and male mice that were bred in-house. Animals were housed in a temperature- and humidity-controlled vivarium on a 12/12 light-dark cycle (lights on at 7 a.m.), and all handling and behavioral testing took place during the light phase. All experimental procedures were approved by the Icahn School of Medicine at Mount Sinai IACUC.

### SURGERY

For all surgeries, anesthesia was induced with 5% isoflurane and subsequently kept at 1-2%. Body temperature was maintained during surgery/recovery with a heating pad below the animal, and ophthalmic ointment was applied to lubricate the eyes. All surgeries followed aseptic surgical technique. Following surgery, animals were given 20 mg/kg ampicillin and 5 mg/kg carprofen (s.c.) per day for 7 days and body weight and general disposition were monitored daily. Animals that underwent gradient refractive index (GRIN) lens implantation were additionally treated with 0.2 mg/kg dexamethasone (s.c.) for 7 days following surgery. All viruses were infused through a glass micropipette at 2 nL/sec using a Nanoject III (Drummond Scientific). Following viral infusion, an additional 5-10 minutes were allowed to pass before retracting the micropipette. Micropipettes were slowly retracted at 0.1 mm/min over 3-5 minutes before being retracted at a faster rate. All coordinates listed below are relative to bregma.

#### Calcium imaging

300 nL of AAV1-syn-flex-GCaMP6f-WPRE was infused into the AHN (AP: −0.5, ML: 0.5, DV: −5.2). A 0.6 mm diameter needle was then slowly lowered to 0.2 mm above the injection site and then retracted in order to create space for the GRIN lens (needle lowered 3x). Lastly, a GRIN lens (0.6 x 7.3 mm, Inscopix, 1050-004597) was lowered to the same coordinates and affixed to the scull using super glue and dental cement. The lens was subsequently covered with Kwik-Sil (World Precision Instruments) followed by a thin layer of dental cement in order to protect the lens prior to baseplating. 4-6 weeks later, while animals were under anesthesia, the lens was uncovered and a baseplate connected to a UCLA Miniscope V4.4 (OpenEphys) was lowered toward the lens until the optimal field of view was identified. The baseplate was then affixed to the skull with super glue and dental cement. The Miniscope was then detached and a dust cover was connected to the baseplate to protect the lens when not imaging.

#### Optogenetics

For optogenetic silencing of AHN neurons, 150 nL of AAV5-hSyn-SIO-stGtACR1-FusionRed (2.1 x 10^13 gc/ml), or AAV5-hSyn-DIO-mCherry (1.1 x 10^12 gc/ml), was infused at the following coordinates at a 20 degree angle: AP: −0.7, ML: 0.5, DV: −5.5. For pan-neuronal silencing, 150 nL of AAV9-hSyn-Cre (1 x 10^12 GC/ML) was co-infused. Subsequently, optic fibers (200 um diameter, 0.5 NA, RWD Life Science) were lowered to 0.6 mm above the injection site, also at a 20 degree angle. Optic fibers were affixed to the skull with super glue and dental cement.

#### Chemogenetics

For chemogenetic activation of AHN neurons, 100 nL of AAV5-hsyn-DIO-HM3Dq-mCherry (2.5 x 10^13 GC/ML), or AAV5-hSyn-DIO-mCherry (4.5 x 10^12^ GC/ML), was infused at the following coordinates at a 20 degree angle: AP: −0.5, ML: 0.5, DV: −5.5. For chemogenetic silencing of inputs to the AHN, a 100 nL mixture of rgAAV-efla-Cre (7.3 ×10^12 GC/ML) and AAV8-hSyn-EGFP (1.5×10^12 GC/ML) was infused at the following coordinates: AP: −0.7, ML: 0.5, DV: −5.5. Separately, 150 nL of AAV5-hSyn-DIO-hM4Di (2.4 x10^13 GC/ML), or AAV5-hSyn-DIO-mCherry (7.3 x 10^12 GC/ML), were infused into either the amygdala (AP: −1.4, ML: 3.1, DV: −5.2), the ventral hippocampus (AP: −3, ML: 3.2, DV: −4.5), or the lateral septum (AP: 0.8, ML: 0.4, DV: −3.3).

#### Retrograde tracing

75 nL of rgAAV-hSyn-EGFP (titer: 1.36 x 10^13 GC/ML) was infused unilaterally at the following coordinates: AP: −0.5, ML: 0.5, DV: −5.5.

### BEHAVIORAL TESTING

#### Single-housing and habituation

All animals were singly housed 1-2 weeks prior to beginning behavioral testing. During this time, they were handled in the vivarium for 1 min/day for 3-5 days. Additionally, animals were habituated to transport to the laboratory for 3-5 days, where they were also handled.

Animals in chemogenetic experiments were habituated to restraint 3x during this time. Animals in Miniscope and optogenetic experiments were habituated to connection to the Miniscope/patch cords ∼5 times. During each of these habituation sessions, animals were connected to the Miniscope/patch cords and returned to their home cage while still connected for 3-5 minutes.

#### Experimental contexts

Stressor 1 (foot-shock stress) and Stressor 2 (auditory stressor), as well as their respective recall sessions, took place in unique experimental contexts, consisting of highly distinct visual, olfactory, auditory, and spatial cues. For Stressor 1, animals were transported from the vivarium in their cages on a cart to the experimental testing room, which was well lit and had an air filter providing ambient sound. Animals were then placed in a brightly lit experimental testing chamber with a grid floor (Med Associates) scented with 5% Simple Green solution. For Stressor 2, with the exception of the iDISCO experiment (see details below), animals were transported from the vivarium in P1000 pipet boxes and carried in a dark cardboard box to the experimental testing room, which was dark except for a dim red light.

Animals were then placed inside of a dark testing chamber (Med Associates) with a flat plexiglass floor and a curved rear wall. The chamber was scented with 1% acetic acid solution.

#### Stressor 1 and Stressor 1 Recall

After a 5 min period of baseline exploration, animals received 10, 1 sec, 1 mA, scrambled foot-shocks, with an inter-shock interval of 30 sec. Animals were taken out of the testing chamber 30 sec after the last shock. For the iDISCO experiment, animals received the same number of shocks, pseudo-randomly distributed over a 60 minute session. When optogenetic inhibition was applied during Stressor 1, blue laser light (473 nm, 5 mW, 20 hz, 20 ms pulse) was continuously administered beginning 30 sec before the first shock and continuing until the end of the session. For Stressor 1 Recall, animals were transported to the same experimental testing chamber for an 8 min test session. When optogenetic inhibition was applied during Stressor 1 Recall, light was administered in 2 min intervals (alternating off-on-off-on). Light-off and light-on epochs were collapsed after similar patterns were observed across the first and second off-on cycles. Stressor 1 and Stressor 1 Recall were separated by at least 2 days.

#### Stressor 2 and Stressor 2 Recall

After a 3 min baseline period, animals were exposed to a single loud auditory stimulus (3 sec, 125-130 dB white noise, 0 ms rise time) that was delivered by a speaker attached to the chamber wall. Animals were removed 10 sec later and returned to the vivarium. When optogenetic inhibition was applied during Stressor 2, blue laser light (473 nm, 5 mW, 20 hz, 20 ms pulse) was continuously administered throughout the entirety of the session. For the iDISCO experiment, Stressor 2 was administered in the home cage in order to avoid cFos evoked by transport habituation. For calcium imaging, a series of 5 auditory stimuli were presented, enabling us to define whether cells reliably fire to the auditory stimuli. In these sessions, after the baseline period was shorted to 2 minutes and the interstimulus interval was also 2 minutes. For Stressor 2 Recall, animals were transported to the same experimental testing chamber for an 8 min test session. Stressor 2 and Stressor 2 Recall were separated by 1 day.

#### Shock sensitivity assay

To assess responses to multiple amplitude shocks, animals were placed in a perceptually distinct conditioning chamber and received a series of 12, 2 second, foot-shocks (6 of each amplitude: 0.25 mA and 1 mA). The first shock occurred 120 seconds after being placed in the chamber and each shock was separated by 60 seconds. Shock amplitudes were presented in a random order with the constraint that a single amplitude was never presented more than twice in a row.

#### Light-Dark Test

The light-dark test was conducted using two interconnected square compartments with an open top (each compartment measured 7.5 in width x 11.25 in height), separated by a 1.5 in wide passageway that could be closed with an opaque sliding divider. One chamber was made of all white acrylic, while the walls of the other were covered in matte black wallpaper and had a red acrylic floor. Overhead lighting provided luminance of 50 lux on the light side. After a 1 minute baseline period in which animals were confined to the dark side, the central divider was raised and the animals could freely explore both sides of the light-dark box. For optogenetic experiments, this period was 8 minutes, and light was administered in 2 minute intervals (alternating off-on-off-on). Light-off and light-on epochs were collapsed after similar patterns were observed across the first and second off-on cycles. For all other experiments, animals were allowed 5 minutes to explore both sides of the chamber. When animals were tested more than once, test sessions were separated by 2 days.

#### Behavior quantification

For analysis of freezing and motion in conditioning chambers when no fibers/cables were attached to the mice, Med Associates Video Freeze software was used (*42*). For measuring freezing and motion in chambers when cables were attached to the animals, as well as in the light-dark test, ezTrack was used (*43*).

### CHEMOGENETIC AND OPTOGENETIC ACTUATION

Actuation of HM4Di and HM3Dq was achieved through intraperitoneal administration of 3 mg/kg cno-dihydrochloride (Tocris), 30-45 minutes prior to behavior, at a volume of 10 mL/kg (dissolved in saline).

For stGtACR1-mediated inhibition, laser light was delivered with a 473 nM laser (OptoEngine LLC) connected via a patch cord to a bifurcating rotary joint (Doric). Consistent with prior reports inhibiting AHN neurons with stGtACR1 (*30*), we utilized 20hz, 20 ms pulse width, 5 mW, illumination. Light intensity was measured from the fiber tip using a light meter (PM100D with S130C attachment, ThorLabs).

### HISTOLOGY

For confirmation of viral placement, GRIN lens placement, and fiberoptic placement, animals were transcardially perfused with 10 mL PBS followed by 10 mL 4% PFA. Brains were then extracted, stored in 4% PFA overnight at 4C, and then transferred to a 30% sucrose/PBS solution before being sectioned at 50 um coronal sections on a cryostat and mounted on slides. Tissue was then imaged on a Leica DM6 epifluorescent microscope. Viral expression and cannula placement was evaluated using the mouse brain atlas of Franklin and Paxinos (*40*).

### iDISCO IMMUNOSTAINING

Following perfusion with 10 mL of 1X PBS and 10 mL of 4% PFA, brains were extracted, any remaining dura and vasculature was carefully dissected off the brain with fine tweezers, and brains were stored overnight in 4% PFA at 4C. Brains were then washed 3x in PBS prior to the iDISCO+ procedure (*16*). All subsequent steps were performed in 5 mL black Eppendorf tubes (fully filled to reduce air). Except where stated, all steps were done while being rotated at 10 RPM.

The general outline of staining/clearing is as follows:

*Day 1:* Tissue was dehydrated in escalating concentrations of methanol (MeOH) in H20, for 1 hr each, at room temperature (20%, 40%, 60%, 80%, 100%). After an additional 2 hrs in 100% MeOH, brains were chilled on ice for 10 min and then incubated overnight in a solution of 66.5% dichloromethane (DCM) and 33.5% MeOH at room temperature.

*Day 2:* Tissue washed twice in 100% MeOH, 3 hours each wash, then chilled at 4C for ∼10min. Subsequently, brains were incubated in chilled 5% hydrogen peroxide (H2O2; 1 volume of 30% H2O2 to 5 volumes MeOH) and kept at 4C overnight, without rotation.

*Day 3:* After allowing samples to warm to room temperature, tissue was rehydrated in decreasing concentrations of MeOH in H20 (60%, 40%, 20%), 1hr each, followed by 1hr in PBS. Subsequently, tissue was incubated 2x in PTx.2 solution (0.2% Triton X-100 in PBS), 1 hr each wash. Lastly, tissue was permeabilized across 2 days at 37C in the following solution: 80% PTx.2, 20% DMSO, 22% glycine (w/v).

*Day 5:* Tissue was transferred to blocking solution (84% PTx.2, 6% donkey serum, 10mL DMSO) and incubated across 2 days at 37C.

*Day 7:* Tissue was transferred to primary antibody (1:2000 rabbit anti-cFos; Abcam 190289) in PTwH (0.2% Tween-20, 0.1% of 10mg/mL heparin solution, in PBS) plus 3% donkey serum and incubated for 1wk at 37C.

*Days 14-15:* Tissue was washed 4-5 times in PTwH at room temperature across 2 days.

*Day 16:* Tissue was incubated in secondary antibody (1: 1000 donkey anti-rabbit IgG AlexaFluor 790; Invitrogen A11374) in PTwH plus 3% donkey serum for 7 days at 37C.

*Days 23-24:* Tissue was washed 4-5 times in PTwH at RT across 2 days.

*Day 25:* Tissue was dehydrated in escalating concentrations of MeOH in H20, for 1 hr each, at RT (20%, 40%, 60%, 80%, 100%).

*Day 26:* Tissue was incubated for 3 hours in 66.5% DCM and 33.5% MeOH at room temperature. After 2 washes in 100% DCM, each 15min, tissue was transferred to 100% dibenzyl ether (DBE).

### LIGHT-SHEET IMAGE ACQUISITION AND QUANTIFICATION

Cleared brains were imaged using a LaVision UltraMicroscope II light sheet microscope using a 1.3x objective lens coupled with 488nm and 785nm lasers for imaging autofluorescence and cFos, respectively. 12-bit horizontal brain images were collected using 3.89uM NA light sheet width in 4.5uM step sizes, spanning from the dorsal-most to ventral-most portion of the brain.

Light sheet images were processed using custom Python-based scripts written in-house to identify cFos cell positions (github.com/ZachPenn/ClearMap2; *zmaster* branch). Images for each animal were preprocessed as follows: 2D images were first converted to 3D arrays, permitting 3D kernel operations. Images were then smoothed using a small median kernel, followed by subtraction of local background fluorescence, estimated using a morphological opening. Small intensity pixel fluctuations were then thresholded and set to zero. To identify cFos puncta in preprocessed images, a distance transform was then calculated, followed by identification of local maxima, corresponding to cell centroids. Importantly, the same image processing parameters were applied to every subject and parameters were selected based upon visual inspection across multiple regions. Images and cell-positions were then aligned to the Allen Brain Atlas (*44*) using the ClearMap2 software package (*16, 45*).

### MINISCOPE IMAGING AND CELL EXTRACTION

Calcium imaging was performed using the UCLA Miniscope, V4.4 (*27*). Parts were obtained from Open Ephys and subsequently assembled in-house. Following baseplating, animals were habituated to wearing the Miniscope over the course of several days while in their home cage. Animals were lightly restrained (held in cupped hand), the dust cover was removed from the baseplate, and the Miniscope was attached to the baseplate and locked into position with a set screw. They were then placed back in their home cage for several minutes with the Miniscope attached and turned on. During this time, the optimal focal plane was identified and this focal plane was held constant throughout the duration of experimental testing by maintaining the setting of the electrowetting lens. Additionally, on each day, the focal plane and field of view was manually compared to the prior day. Miniscope videos were acquired at 30 frames per second.

Calcium imaging videos were processed using the open-source software, *minian* (*46*). In brief, for each session, videos were down-sampled to 15 fps and corrected for translational motion. To improve signal quality, a minimum projection for each video was then subtracted from each frame to remove vignetting, a median filter was applied to remove granular noise, and a morphological opening was performed to identify – and subsequently filter out – local fluctuations in background. To identify potential cell locations, local maxima in the field of view were defined across the video. These local maxima were then utilized as input to the CNMF algorithm. CNMF parameters were chosen for each animal based upon the auditory stressor session in a visually guided manner using *minian*. These parameters were then applied to every video for that animal. Parameters were selected blind to group membership. For all analyses, we utilized the raw signal for each cell, obtained by multiplying the spatial footprint resulting from CNMF by its temporal activity (i.e., each pixel has a weight for each cell, and the weighted average of fluorescent intensity values across time for these pixels comprise a cell’s activity). Cross-registration of cells across sessions was performed using *minian*.

### STATISTICAL ANALYSIS

All statistical analysis were performed in R. All data and statistical analysis are available at github.com/ZachPenn/AHN. Group sizes are listed in each figure legend.

#### Whole Brain cFos Analysis

Within the Allen Mouse Brain Atlas (*44*), regions are classified in a nested hierarchy (e.g., infralimbic layers 1 and 2 are nested within the infralimbic cortex which is in turn nested within the isocortex). For all cFos analyses, the lowest available level of this hierarchy was used, providing the most granular regional information. The olfactory bulbs and cerebellum were excluded from all analyses due to frequent damage resulting from dissections.

Additionally, any region that had very low cFos counts across groups was excluded (median counts per group all less than 1). This left a total of 454 brain regions across which subsequent analyses were conducted.

##### Group differences in regional cFos counts

To compare cFos counts across groups, negative binomial regression was performed using the MASS package in R (*47*). For each region, count data was modeled as a function of group (y=B_0_+B_1_*NT+B_2_*T+e), with the three groups consisting of those animals that did not receive Stressor 2 (Control, Ctrl), those that did not receive Stressor 1 but received Stressor 2 (NS), and those that received both stressors (S). We first assessed whether group membership was a significant predictor of cFos counts by comparing the full model listed above to an intercept only model (y=B_0_+e). Amongst the 90 regions that displayed group variation in cFos counts following FDR correction, we then assessed whether S and NS groups displayed differential cFos counts using post-hoc contrasts with the *multcomp* package (*48*).

For this contrast, we display regions which surpass FDR correction, as well as those regions that display differential activation at an uncorrected threshold of p=0.05 in order to better capture the general pattern of results. The results of statistical tests for each region can be found in Supplementary Table 1. Additionally, summary statistics for each region/group can be found in Supplementary Table 2. For full count information and analysis scripts, please see github.com/ZachPenn/AHN.

##### Correlational analysis and hierarchical clustering

For all correlational and hierarchical clustering analyses, cFos counts were z-scored separately for each group. Because analyses focused on NS and S animals, which had a substantial spread of counts and moderate sample size (n=10/group), Pearson correlations were used as the bases for these analyses.

For hierarchical clustering, the region-region correlation matrix relating counts between brain regions (R) was first converted to an adjacency matrix (A = 1-R), and subsequently to a set of distances using the *stats* package in R (*49*). Agglomerative hierarchical clustering was then performed on these distances using the average distance between clusters. Cluster thresholds were set using the ‘elbow’ method, where cluster number is increased until within-cluster variance no longer decreases substantially when increasing cluster number (Fig S3). Additionally, we attempted to keep within-cluster correlations high (R>=0.5).

To compare cluster ‘strength’ across groups, we initially performed hierarchical clustering on animals exposed to Stressor 1 (S), as described above. After identifying the cluster containing the AHN in these animals, we examined the correlation matrix exclusively for regions within this cluster in S and NS animals. In particular, we extracted the mean off-diagonal correlation, as this reflects overall within-cluster relatedness, and compared this value across groups (i.e., avg(R_t_) – avg(R_NT_)). This was compared to a null distribution computed by shuffling the off-diagonal correlation values across groups and computing the same difference of mean correlations 1000 times. A similar approach was taken to compare the overall correlation matrix from each group.

To compare the number of correlations between the AHN and other brain regions between animals that received Stressor 1 and those that did not, we compared the number of significant correlations between groups using the chi-square test. Of note, while we used uncorrected significance (p=0.05) to more broadly illustrate functional connectivity of the AHN, using FDR correction provides a similar result (Chi-sqaure contingency test of S vs NS: χ^2^=10.2, p<0.01).

#### Calcium Imaging Analysis

For each cell and session, activity was first normalized by computing a z-score and then subtracting the minimum value from across the entire session, such that each cell’s activity was in standard deviation units with a minimum value of 0. For plotting cell responses, activity was first averaged in 200 ms bins. For comparing responses of cells to various amplitude shocks, activity was binned into 1 second intervals and cells were treated as individual datapoints.

##### Classification of responsiveness

To classify whether a cell was responsive to a particular stimulus (e.g., shock), the average change in activity in the 5 seconds following stimulus onset relative to the 5 seconds preceding stimulus onset was first assessed. A null distribution was then obtained by randomly shuffling stimulus onset times and recalculating this average difference 1000 times. Stimuli that had a response above the 95^th^ percentile of this null distribution were considered responsive.

#### Behavioral analysis

For analysis of behavioral data, omnibus ANOVA were conducted using the package ezANOVA with type 3 degrees of freedom. The white adjustment was implemented to correct for heterogeneity of variance using heteroscedasticity corrected standard errors (‘hc3’). For repeated measures ANOVA, the Greenhouse-Geisser correction was implemented when the assumption of sphericity was not met. F and t values are rounded to the nearest tenth and hundredth, respectively. Where F values were less than .1, F is listed as 0.

